# Maxillary constriction causes nasal septum deviation and deformity of the nasal floor

**DOI:** 10.64898/2026.02.17.706297

**Authors:** Mani Alikhani, Eileen Uribe-Querol, Daniel L. Garzón, Chinapa Sangsuwon, Jeanne Nervina, Fanar Abdullah, Mona Alikhani, Nuria Galindo-Solano, Janeth Serrano-Bello, Lucia Pérez-Sánchez, Guillermo Villagómez-Olea, Francisco J. Marichi-Rodríguez, Cristina C. Teixeira

## Abstract

**Introduction:** We investigated the direct effect of transverse maxillary constriction on nasal septal deviation (NSD) and nasal floor slanting.

**Methods and Materials:** 60 growing Wistar rats (21days old) were divided into four groups (n=15): 1) Experimental Group 1 received active constriction force (100cN), 2) Experimental Group 2 received active expansion force (100cN), 3) Sham received the same spring as Experimental Groups without receiving any active force, and 4) Control group did not receive any appliance. Samples were collected after 28 days for microcomputed tomography (μCT) analysis.

**Results:** Experimental Group 1 demonstrated maxillary constriction (both skeletal and dental), accompanied by mandibular shift on closure, clockwise mandibular rotation, and increased mandibular plane angle and facial height. Constriction also produced severe nasal floor slanting in the molar area that extended posteriorly. Nasal floor canting was accompanied by a slanted vomer and a C-shaped NSD. The direction of nasal floor canting and mandibular shift was always similar. Experimental Group 2, on the other hand, was not associated with nasal deviation, and a slight slanting of the nasal floor was observed only when there was a mandibular shift.

**Conclusion:** Our study suggests that the constricting transverse forces applied to the maxilla can induce slanting of the nasal floor and, consequently, the vomer, which in turn can lead to nasal septal deviation. Slanting of the nasal floor is caused mainly by rotation of the hemimaxilla in response to transverse forces and changes in occlusal forces due to a mandibular shift.

## Introduction

Why we can have a nasal septal deviation is not clear. Acquired and congenital factors that contribute to the development of nasal septal deviation, such as trauma and craniofacial anomalies, are well-studied [1–10]. However, these factors alone cannot explain the high incidence of nasal septal deviations. This is important because nasal septal deviation is among the leading causes of nasal obstruction, along with enlarged adenoids, concha bullosa, and inferior turbinate hypertrophy, all of which may lead to mouth breathing. Chronic mouth breathing over an extended period can increase the constricting forces on the maxilla, thereby affecting its transverse and vertical growth [11–13]. Further maxillary constriction can cause occlusal disharmony, narrowing of the pharyngeal airway, alterations in tongue posture resulting in retroglossal airway narrowing, and potentially contribute to obstructive sleep apnea [14–19].

Previous studies have shown that nasal septal deviation is rare among long-snouted mammals and is more common in mammals with shorter faces [3, 20]. Similarly, from an evolutionary point of view, it has been suggested that humans may have developed a high incidence of nasal septal deviation as a result of the continued diminution of our face [20, 21]. It has also been reported that individuals with a more retrognathic nasomaxillary complex exhibit a greater magnitude of septal deviation compared to individuals with a more mesognathic midface [9] . In addition, nasal septal deviation is more common in individuals with reduced facial height [22]. Similarly, experimental reduction of the maxillary length through surgery restricts normal septal growth and induces septal deviation [23]. These observations suggest a possible developmental etiology for nasal septal deviation, possibly due to a constricted maxilla. Based on this theory, deviations occur when cartilage volume exceeds the volume of available space for the cartilage due to a decrease in the size of the maxilla in the transverse dimension. This suggests reciprocal effects between the nasal septum and maxilla; a constricted maxilla produces nasal septal deviation, and in turn, nasal septal deviation through mouth breathing worsens the maxillary constriction.

However, studies investigating maxillary transverse deficiency as a developmental factor in nasal septal deviation are rare and indirect, focusing mainly on identifying an association between the incidence of nasal septal deviation and the incidence of maxillary constriction [24–27].

Here, for the first time, we investigate the direct effect of maxillary constriction on the development of nasal septum deviation in growing rats. Furthermore, we investigate the mechanism by which maxillary constriction may contribute to nasal septal deviation. Considering that the incidence of maxillary constriction is increasing in humans[3, 28–37], establishing a relationship between maxillary constriction and the development of nasal septal deviation is a significant health concern and justifies preventive measures, as maxillary constriction can affect the shape of the nasal septum. Therefore, early treatment of maxillary deficiency with orthopedic tools can have significant health benefits.

## Methods and Materials

### Animal Study

Growing Wistar rats (n = 60, average weight 47g, 21 days old) were studied at two centers: New York University and the Universidad Nacional Autónoma de México (UNAM). Animals were treated in accordance with the protocol approved by the New York University Institutional Animal Care and Use Committee and the Guidelines of the Mexican Law of Animal Protection (NOM-062-ZOO-1999). All experiments were approved by the local Institutional Animal Care and Research Advisory Committee (CICUAL, ID 10389), from Universidad Nacional Autónoma de México (UNAM), and New York University, and under International Laws of ethical care and use of animals (National Research Council (U.S.) et al., 2011) and the National Research Council’s Guide for the Care and Use of Laboratory Animals (IACUC Protocol # 151003).

Rats at this stage are in a growing phase, and their molars are fully erupted. All animals were housed in polycarbonate cages in a 12-hour light/dark environment at a constant temperature of 23°C and fed a specially prepared soft diet with tap water *ad libitum*. The mixture consisted of 200 g of standard chow (Teklad, Envigo), 600 mL of water, and 45 g of gelatin, which was homogenized and subsequently molded into bars to facilitate consumption by the rats. This diet was used to ensure adequate nutritional intake while minimizing the possibility of dislodging the appliance. Animals were randomly divided into four groups: 1) an untreated Control, 2) a Sham Group, 3) Experimental Group 1 (Constriction), and 4) Experimental Group 2 (Expansion).

### Transverse force application

On day 0, animals from the Sham and Experimental Groups were sedated with isoflurane delivered through the SomnoSuite anesthesia system (Kent Scientific). Isoflurane was administered at 1% during the initial sedation phase, increased to 3% to achieve deep anesthesia, and subsequently reduced to 0.5% for maintenance. Anesthesia was verified by lack of response to toe-pinch. Springs were fabricated from 0.016” stainless steel wires (3M Unitek, Monrovia, CA, USA). The Experimental Groups received a calibrated, custom-designed constriction or expansion spring that delivered compression or tensile transverse force (100cN) to the molars (Figure 1). This force was selected based on previous studies demonstrating that high cellular activity was observed in the suture at 100cN [38]. Considering that rats would have a normal vertical chewing force, averaging between 54 and 75 N, this force is not regarded as excessive [39]. The appliances were cemented with composite resin on the interproximal and occlusal surfaces of the molars. The Sham group received a similar spring, but it was not activated and produced no forces. Compressive or tensile forces were applied for 28 days. Animals were monitored daily under inhalational anesthesia with isoflurane–nitrous oxide as the standard method of general anesthesia to ensure the integrity of the appliance and for the need to reinstall due to masticatory forces. No significant differences were observed among groups.

**Figure 1:**
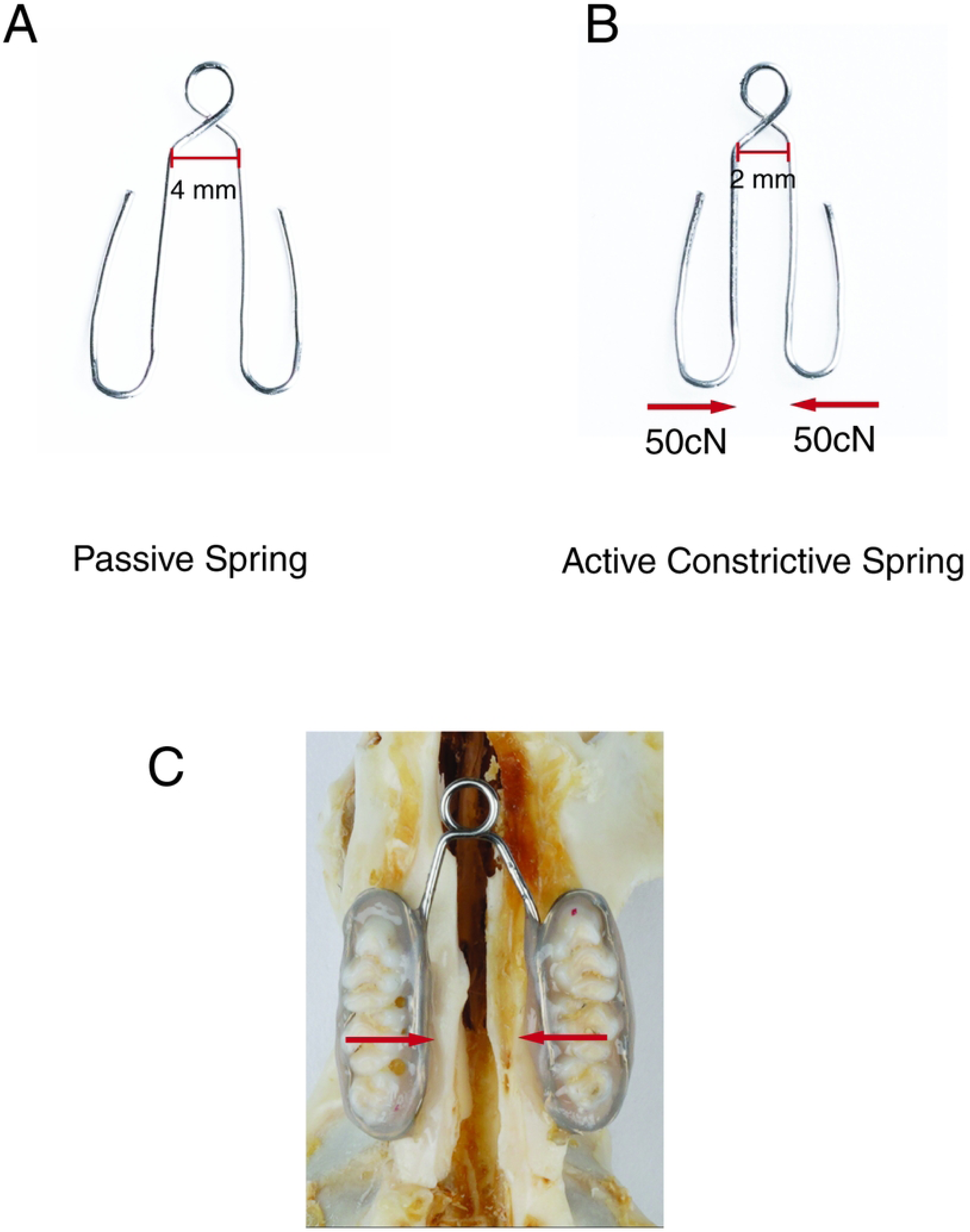
Calibrated spring used to produce transverse forces when installed on the maxillary molars. (**A)** Springs were fabricated from 0.016” stainless steel wires (3M Unitek, Monrovia, CA, USA). (**B)** For Experimental Group 1, the springs were calibrated using a digital force gauge to produce 100cN compression force upon insertion (50cN per side). For Experimental Group 2, springs with a similar design, except the springs were expanded by 2 mm, were used to make a 100cN expansion force (not shown). (**C**) Photograph of rat maxilla with a compression spring held in place on the molar teeth by the application of the light-cured composite.

### Micro-CT Imaging

Animals were euthanized by CO_2_ narcosis on day 28, and samples were collected for micro-CT analysis (μCT). After euthanasia, the whole skull was dissected and fixed for over 72 hours with 4% (w/v) paraformaldehyde in 0.1 M phosphate buffer, pH 7.4, followed by storage in 70% ethanol.

The whole maxilla was scanned by micro-computed tomography (µCT; Skyscan1172; Bruker microCT, Kontich, Belgium). The specimens were scanned at 13.55µm voxel size, 100 KV, 0.300 degrees rotation step (192.30 degrees angular range), and a 1910ms exposure per view in 70% ETOH. Results were analyzed utilizing the NRecon (1.6.9.16) software on the HP open platform (OpenVMS

Alpha Version 1.3-1 session manager) for 3D reconstruction and viewing of images. Superimpositions were performed using Amira 6.0.0.

### Intra-oral and Intra-nasal Measurements

Palatal width (at the level of intersection between alveolar bone and palatal walls) and interdental width (the distance between the height of contour of the first molars) were analyzed in the mid-coronal plane at the level of the upper first molar from the three-dimensional images of µCT (Figure 2A). Changes in the upper molar inclination were studied by measuring the angle (a) formed by the long axes of upper right and left first molars in the same plane.

**Figure 2:**
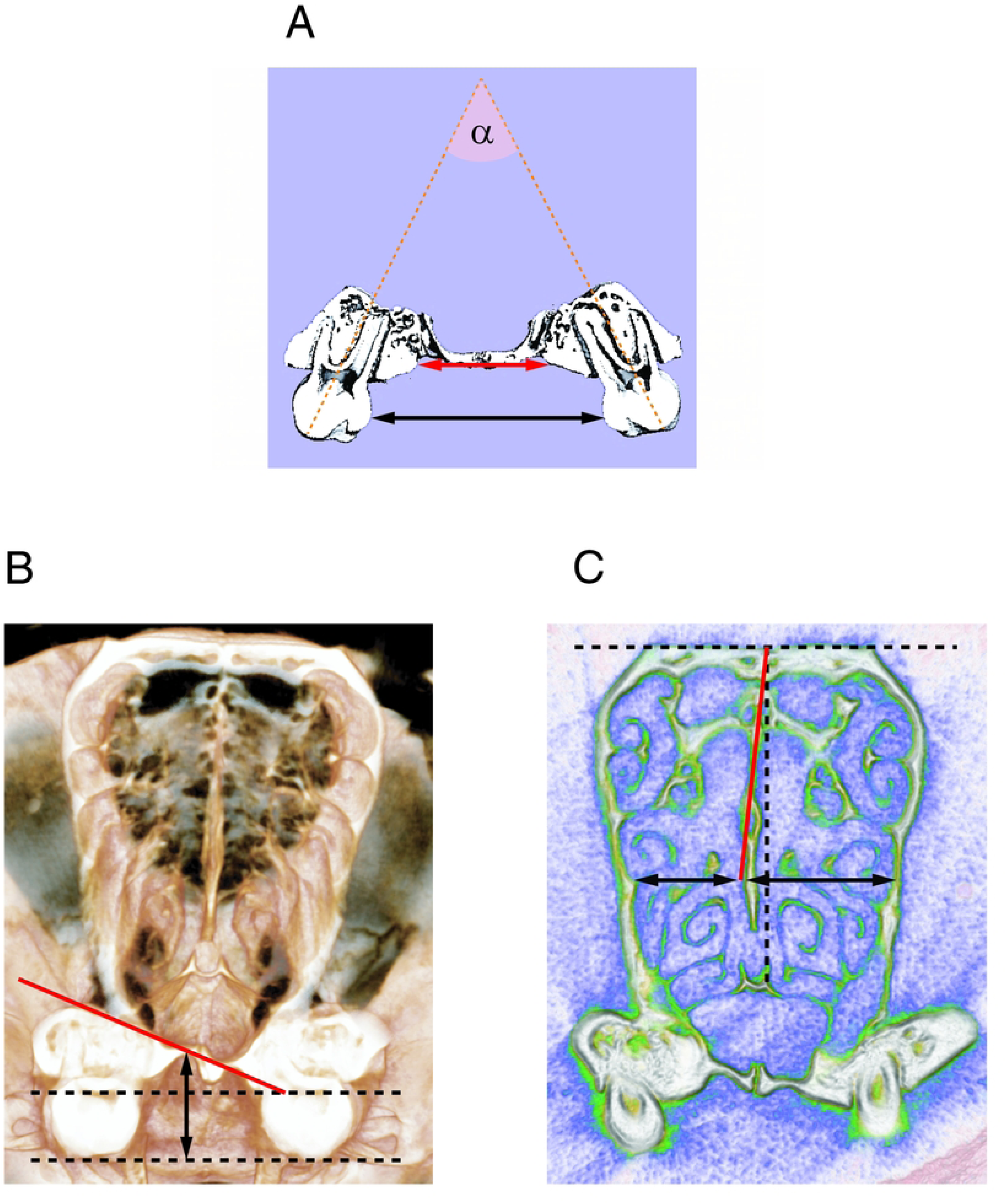
Schematic of Intra-Oral and Intra-Nasal measurements. (**A**) Schematic showing the measurements used to study the effect of constricting forces on the width of the dental arch and palate, and inclination of upper teeth. The width of the palate, as the distance between palatal walls at the level of intersection between alveolar bone and palatal walls (red arrow), and interdental width as the distance between the height of contour of the first molars (black arrow) were analyzed at the level of the mid-coronal plane of the upper first molar from the three-dimensional images of µCT. Changes in the inclination of the upper molars were studied by measuring the angle (a) between the dental long axes. **(B)** The slant of the nasal floor was investigated by measuring the angle a line drawn parallel to the orbits (dashed red line) and another line was drawn along the steepest slant of the nasal floor (blue line) in the coronal section of the µCT images, where the slant was maximum. In the same image, the palatal depth (green arrow) was measured from the highest point of the palate perpendicular to the occlusal plane (black line). **(C)** The degree of the nasal septum deviation was assessed by measuring the angle between the tangent line to the upper part of the nasal septum (dashed yellow line) and a perpendicular line drawn from the horizontal line connecting the center of the two orbits (black lines) on the soft tissue view of the µCT images, where the deviation was maximum. The width of the left and right nasal cavities was measured on the area of maximum deviation on a horizontal line from the septum to the lateral walls of the nasal cavity (red arrows) on the same image.

The slant of the nasal floor was measured on the coronal plane of the µCT image, specifically in the area where the slant was most pronounced. One line was drawn tangent to the nasal floor and parallel to the orbits, and another line was drawn along the steepest slant of the nasal floor. The angle between these lines was measured and recorded as the slant of the nasal floor (Figure 2B). On the same µCT image, the palatal height was measured as the perpendicular distance from the highest point on the palate to the occlusal plane (Figure 2B).

The degree of the nasal septum deviation was assessed by measuring the angle between the tangent line to the upper part of the nasal septum and a perpendicular line drawn from the horizontal line connecting the center of the two orbits. (Figure 2C). The width of the left and right nasal cavities was measured on the area of maximum deviation on a horizontal line from the septum to the lateral walls of the nasal cavity (Figure 2C). Measurements were performed on soft tissue view of the coronal section of microCT images, where the deviation was most significant.

### Extra-oral Measurements

Skull changes in response to constriction were studied by measuring the angle between the mandibular plane (the line tangent to the lower border of the mandible) and the palatal plane (the line tangent to the palate). The anterior facial height was measured on the perpendicular line to the mandibular plane from the most anterior point of the nasal bone (Figure 3A). The posterior facial height was measured from the highest point in the condylar process to the lowest point on the gonial angle (Figure 3B). Maximum condylar width was measured in the coronal view from the medial to the lateral heights of contour (Figure 3C).

**Figure 3:**
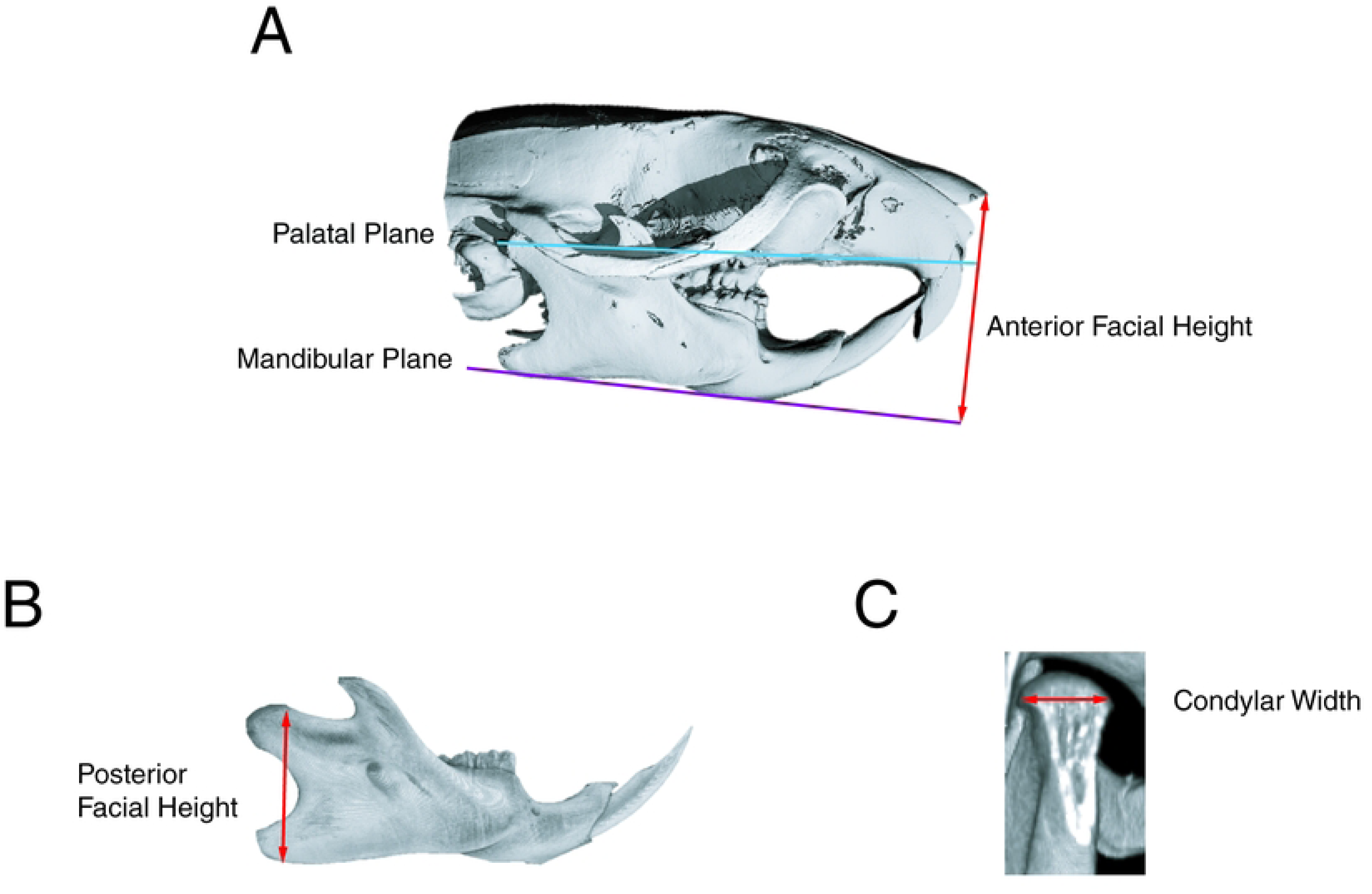
Schematic of Skeletal Measurements. A sagittal image of µCT was used to study the changes in the skull in response to constriction. **(A)** The mandibular plane (yellow line) and palatal plane (red line) were defined as the line tangent to the lower border of the mandible and the line tangent to the palate, respectively. The anterior facial height (black arrow) was measured on the perpendicular line to the mandibular plane from the most anterior point of the nasal bone. **(B)** The posterior facial height (yellow arrow) was measured from the highest point in the condylar process to the lowest point in the gonial angle. **(C)** The condylar width (red arrow) was measured on a coronal section of the condyle from the height of the contour of the condyle on the medial side to the height of the contour of the condyle on the lateral side at the widest point.

### Statistical Analysis

Two examiners completed all morphological quantifications. The random and systematic errors were calculated using the formula described by Dahlberg and Houston [40, 41]. Both the random and systematic errors were found to be small for intra-observer (0.015 and 0.017 mm, respectively) and inter-observer (0.018 and 0.02 mm, respectively).

Significant differences between the Experimental Groups, Shams, and Controls were assessed using analysis of variance (ANOVA). A pairwise multiple-comparison analysis was performed using Tukey’s post hoc test. Two-tailed p-values were calculated; p < 0.05 was set as the level of statistical significance.

## Results

### Constriction significantly decreased the palatal width, interdental width, and dental angulation

Application of constricting force to the maxilla was accompanied by narrowing of the upper dental arch in the area of the molars (Figure 4A). To better understand the nature of this narrowing, the width of the palate, interdental width, and angulation between the long axis of the teeth were analyzed at 28 days at the level of the mid-coronal plane of the upper first molar from the three-dimensional images of µCT as described (Figure 2A).

**Figure 4:**
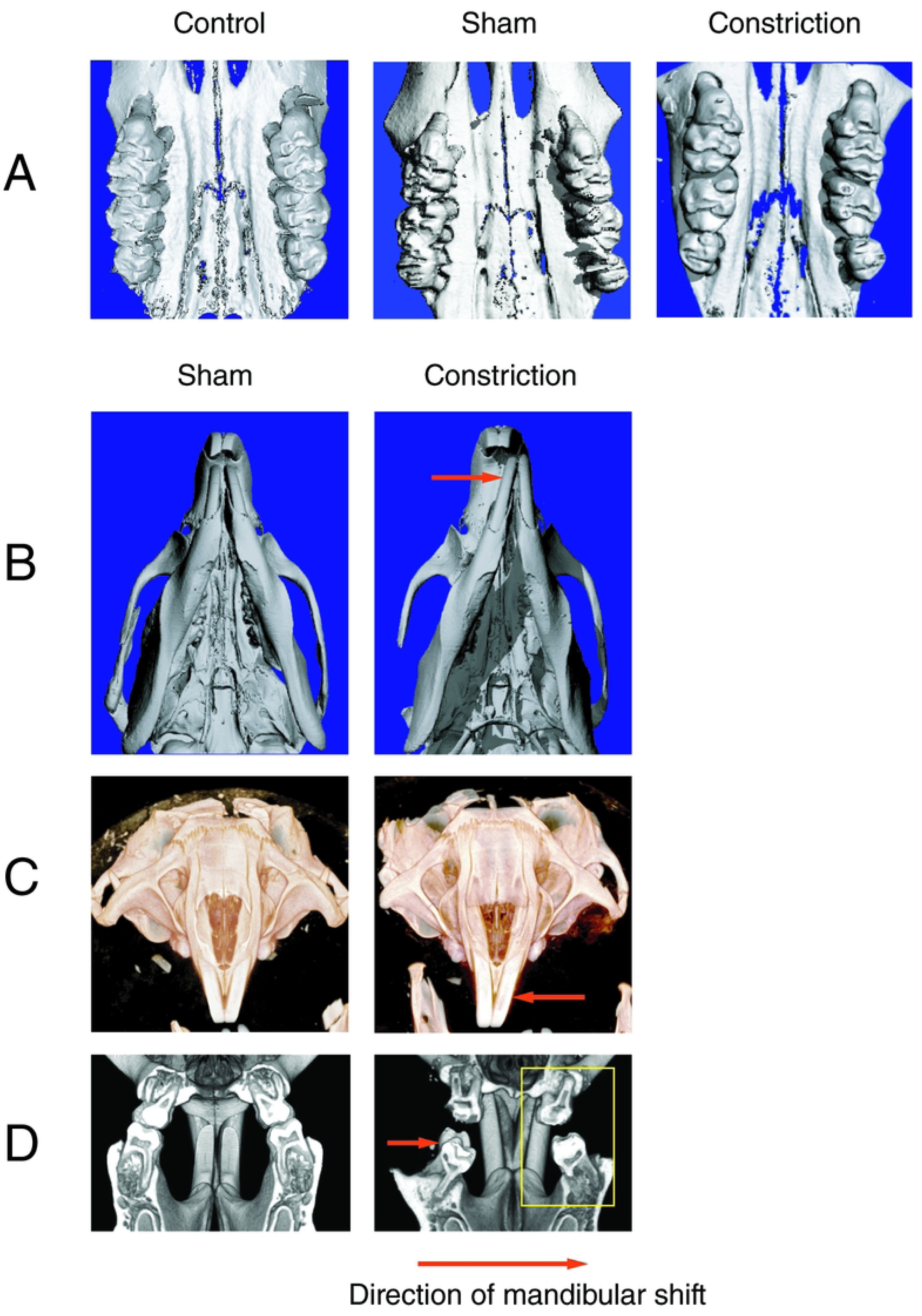
Constriction caused narrowing of the maxilla: **(A)** Occlusal view of µCT scan of the rats that received constricting forces demonstrates a decrease in transverse dimension of the palate and intermolar distance. **(B)** The decreased palatal width was accompanied by a mandibular shift, in most cases toward the left side, and a change in bite between anterior teeth (yellow arrow) and inclination of some of the anterior teeth **(C)**. In addition, the constriction of the maxilla and shift of the mandible produced a posterior crossbite in the non-working side (red circle) and an increase in inclination of the posterior teeth on the working side (white arrow) **(D).** The black arrow demonstrates the direction of mandibular shift.

Constriction significantly decreased dental and palatal width compared with the Control and Sham groups at 28 days, with a statistically significant difference (p<0.01) (Table I). The palatal movement of the teeth and alveolar bone includes significant up-righting of molars as observed with a substantial decrease in the angulation between the long axis of the upper first molars, which was statistically significant (p<0.01). The magnitude of dental changes was greater than that of skeletal changes, as demonstrated by a greater decrease in dental width than in palatal width. Since no difference was observed between the Sham and Control groups, only data from the Sham group is presented in the rest of the article.

**Table I:**
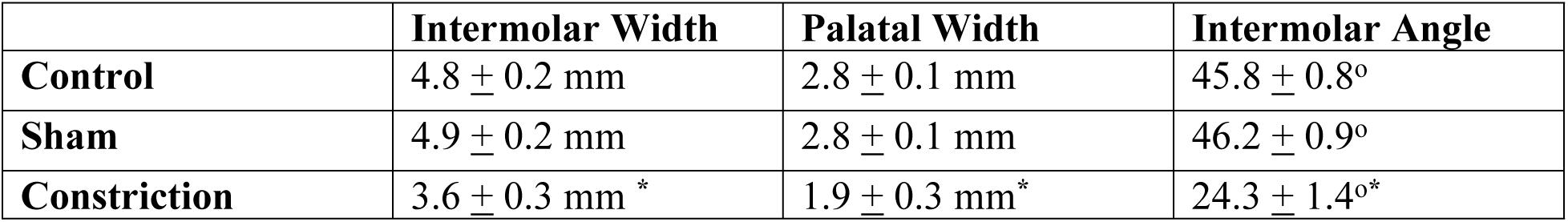
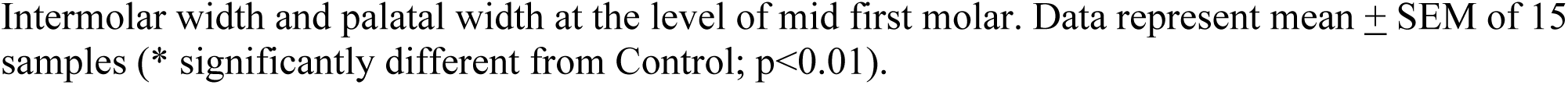
Transverse changes in the maxilla in response to constriction.

|  | **Intermolar Width** | **Palatal Width** | **Intermolar Angle** |
| --- | --- | --- | --- |
| **Control** | $4.8 \pm 0.2$ mm | $2.8 \pm 0.1$ mm | $45.8 \pm 0.8^\circ$ |
| **Sham** | $4.9 \pm 0.2$ mm | $2.8 \pm 0.1$ mm | $46.2 \pm 0.9^\circ$ |
| **Constriction** | $3.6 \pm 0.3$ mm * | $1.9 \pm 0.3$ mm * | $24.3 \pm 1.4^\circ$ * |

Intermolar width and palatal width at the level of mid first molar. Data represent mean ± SEM of 15 samples (\* significantly different from Control; $p < 0.01$ ).

Intermolar width and palatal width at the level of mid first molar. Data represent mean ± SEM of 15 samples (* significantly different from Control; p<0.01).

Maxillary constriction caused the animals to shift their mandibles into a comfort zone for biting, and, in these experiments, nine of the fifteen rats in the constriction group moved to the left side; however, the difference in the direction of shift between animals was not statistically significant (p>0.05). The result of the mandibular shift was an asymmetric bite in the anterior teeth (Figure 4B), leading to changes in the mesio-distal angulation of some of the incisors (Figure 4C). This also produced a posterior crossbite (Figure 4D) with a consequent adaptive change in the inclination of the lower molars, with more inclination of the posterior molars on the non-crossbite side.

### Constriction of the maxilla resulted in clockwise rotation of the mandible

Constriction of the upper jaw and narrowing of the dental arch were accompanied by relative extrusion of maxillary teeth (up-righting of molars in the coronal plane), which caused clockwise rotation of the mandible and an increase in the mandibular plane angle compared to the palatal plane. This was accompanied by an increase in anterior facial height and a decrease in posterior facial height, both of which were statistically significant (p< 0.05) (Figure 5A and B) (Table II). An increase in mandibular plane angle was accompanied by an increase in overjet (the horizontal relation between the upper and lower incisors) and a decrease in overbite (the vertical relation between the upper and lower anterior teeth), resulting in an animal with a Class II appearance (Figure 5A). Change in biomechanics of the jaw was accompanied by condylar adaptation and a decrease in the width of the condyles that was statistically significant (p<0.05) (Figure 5C).

**Figure 5:**
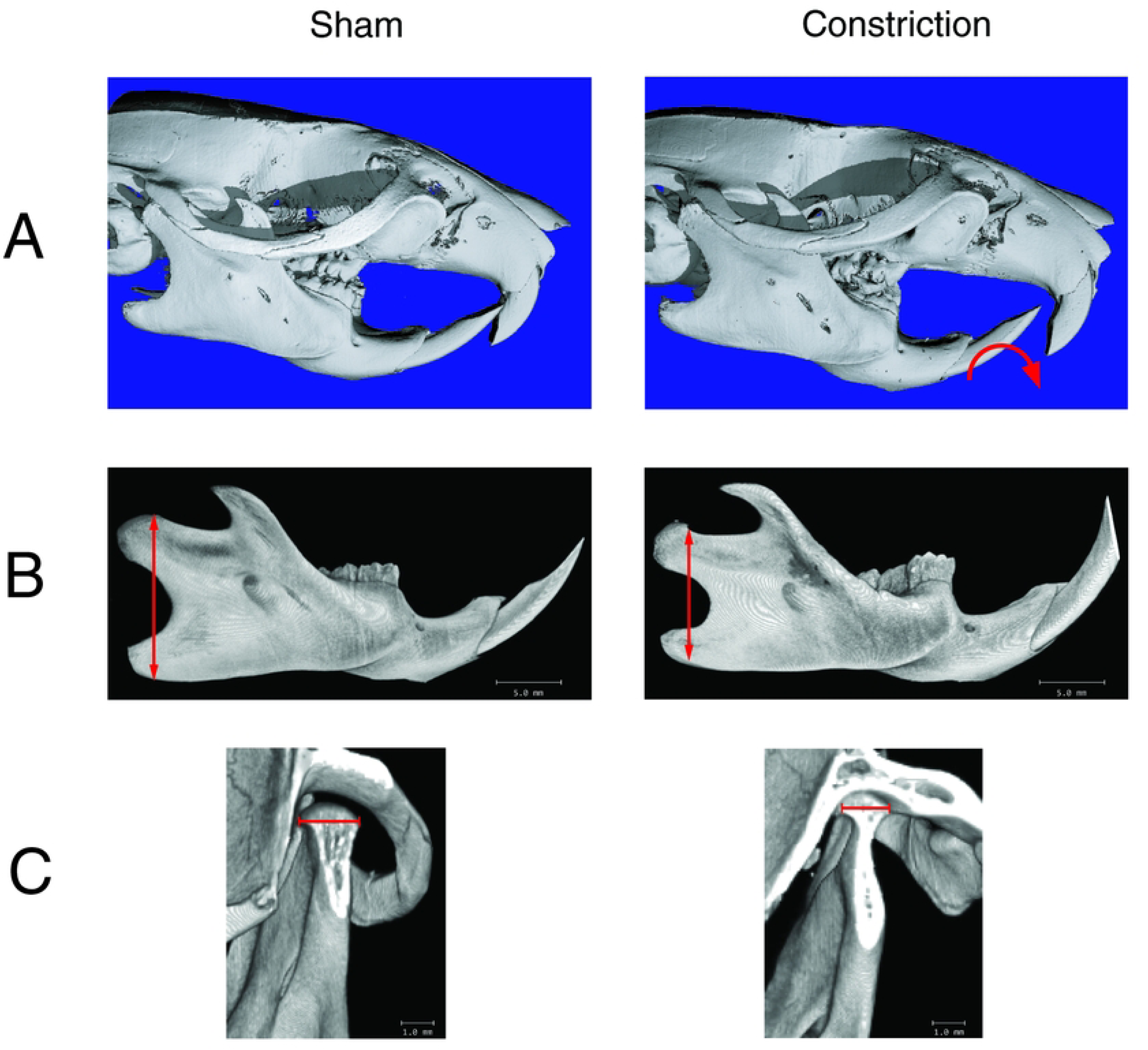
Constriction caused a clockwise autorotation of the mandible and increased facial height. **(A)** A comparison of the sagittal view of the µCT scan of Experimental Group 1, which was exposed to constriction, and the Sham group, demonstrated a clockwise rotation of the mandible following constriction. This gives the skull a slight Class II appearance with an increased overjet and a decreased overbite. In addition, the mandible of the Experimental Group 1 rats demonstrated a reduction in posterior facial height (**B)** and the width of the condyles **(C).**

**Table II:** General skull characteristics after 28 days of constriction.

|  | Angle between the mandibular plane and the palatal plane | Anterior facial height | Posterior facial height | Condylar width |
| --- | --- | --- | --- | --- |
| Sham | $8^{\circ} \pm 1^{\circ}$ | $15.7 \pm 0.8$ mm | $12.4 \pm 0.7$ mm | $1.81 \pm 0.2$ mm |
| Constriction | $15^{\circ} \pm 2^{\circ} *$ | $20.2 \pm 1.3$ mm * | $9.2 \pm 0.8$ mm * | $1.23 \pm 0.3$ mm * |

Data represent mean ± SEM of 15 samples (\* significantly different from Sham $P < 0.05$ )

### Maxillary constriction caused nasal floor slanting

µCT images of Experimental Group 1 rats revealed an average slant of 26° in the nasal floor, which is statistically significant compared to the Sham Group (p< 0.01). None of the Sham or Control Groups demonstrated any nasal floor deformity (Table III).

**Table III:** Nasal floor inclination.

|  | **Nasal Floor Angle** | **Palatal Depth** |
| --- | --- | --- |
| **Sham** | **$0 \pm 1^\circ$** | **$3.1 \pm 0.1 \text{ mm}$** |
| **Constriction** | **$26 \pm 3^\circ$ \*** | **$4.8 \pm 0.5 \text{ mm}^*$** |

Data represent mean ± SEM of 15 samples (\* significantly different from Sham $P < 0.01$ )

In Experimental Group 1, nine animals demonstrated a slant extending from right to left. In comparison, in six animals, the slant was extended from left to right with no statistical difference (p>0.05) (Figure 6A). The direction of slant and the direction of the shift of the mandible were the same.

**Figure 6:**
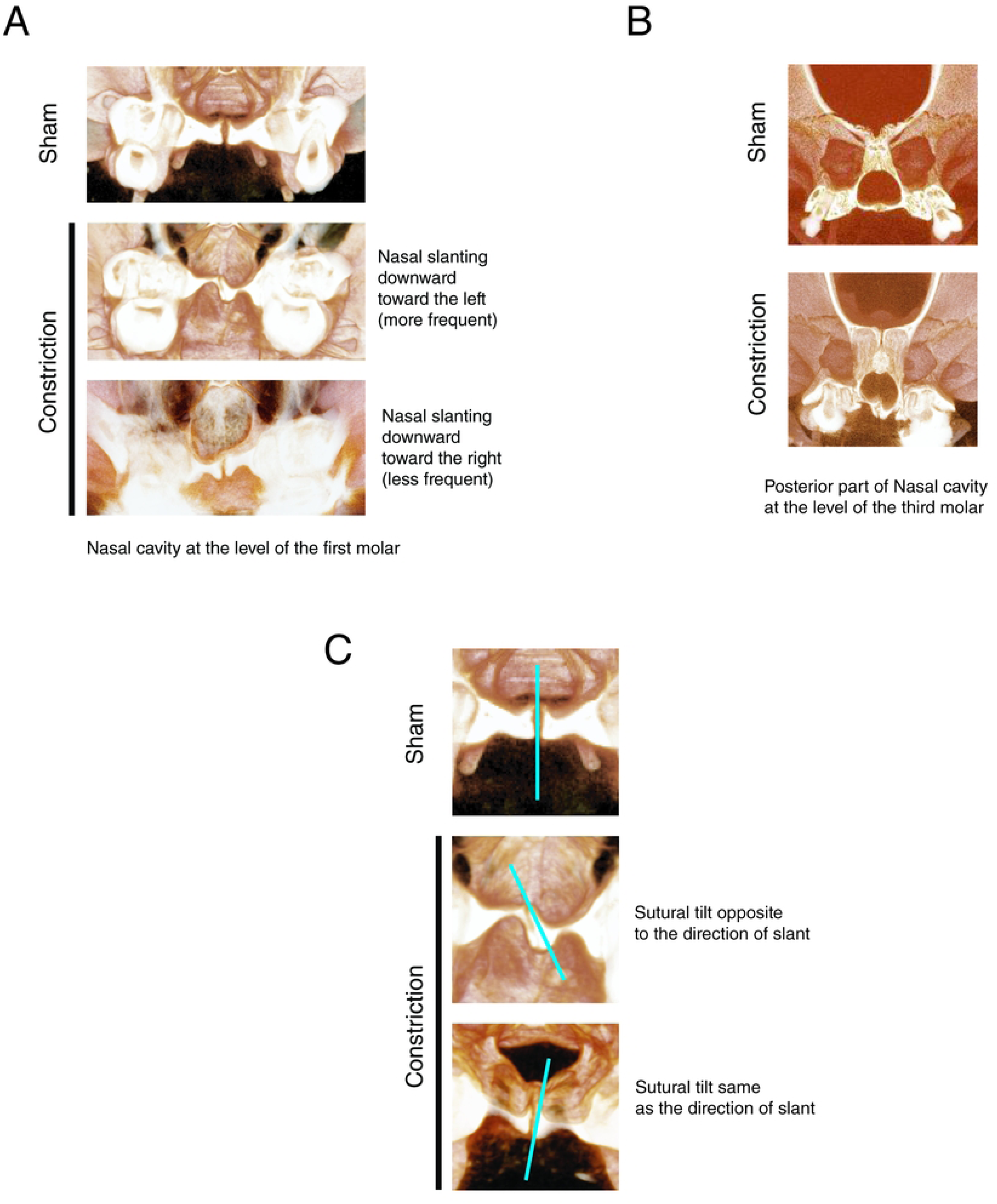
Maxillary constriction caused nasal floor slanting. The dominant slant observed was from right to left; however, in a few animals, the slant extended from left to right **(A)**. The palatal slant followed the same direction as the slant of the nasal floor. **(B)** The nasal floor slant did not include the anterior region of the septum and started in the area of the molars and was extended to the posterior part of the septum**. (C)** Coronal view of the mid-palatal sutures also demonstrated a vertical tilt in one direction (yellow line). The sutural tilt was independent of the direction of the nasal floor slant. The Sham did not show any vertical tilt in the mid-palatal suture. The horizontal position of the suture showed no variability.

The deformity of the nasal floor started in the area of the molars, where the constricting force was applied, but not in the anterior region of the septum. The slant in the nasal floor was continuous, varying in degree, and extended to the posterior nasal cavity (Figure 6B).

The palate followed the nasal floor slanting. The direction of the palatal slant in all animals was the same as that of the nasal floor slant. In addition, a significant increase in palatal depth was observed compared with the Sham group (Table III).

The nasal floor and palatal floor deformities also affected the suture morphology (Figure 6C). The deformity did not change the horizontal location of the mid-palatal suture; however, it changed the coronal direction of the suture, which demonstrated that the coronal direction of the mid-palatal suture can be altered due to constricting forces. The direction of the sutural tilt was independent of the direction of slanting (Figure 6C).

### Maxillary constriction caused a nasal septal deviation

Maxillary constriction was accompanied by curvature of the nasal septum. Nasal septum deviation was limited to the area of the molars; the anterior part of the nasal septum did not demonstrate any deviation. While the most frequent direction of deviation was toward the lowest part of the nasal septum slant, fewer animals demonstrated opposite deviation with no statistical difference (p>0.05) (Figure 7A). The magnitude of deviation from the vertical position in the average was approximately 11°, which was significant compared to the Sham Group (p < 0.03) (Table IV). In our experiments, the deviation was always C-shaped.

**Figure 7:**
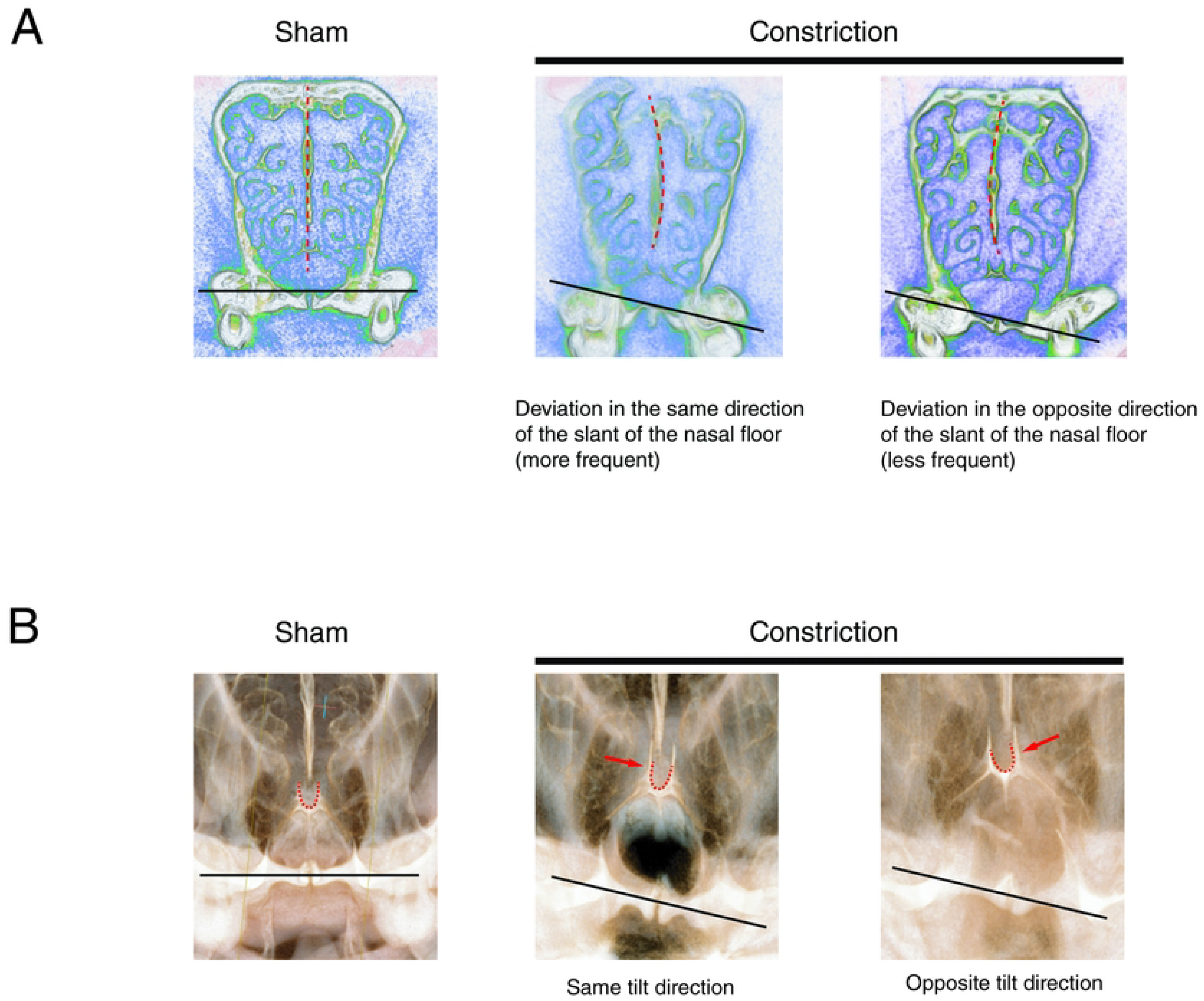
Maxillary constriction was accompanied by nasal septum deviation. **(A)** Deviation of the nasal septum (black dashed line) was observed in response to the application of constricting forces. Sham group septum did not show any deviation. The more frequent direction of deviation was toward the lower part of the nasal floor slant (red arrow). However, in fewer animals, the direction of deviation was opposite and toward the higher part of the nasal floor slant. **(B)** The vomer channel that holds the bottom part of the nasal septal cartilage demonstrates the same tilt as the direction of the septum in all cases. However, compared to the nasal floor, it could adapt either the same or opposite direction (more frequently toward the lower part of the nasal floor slant (yellow arrow). The Sham group did not show any tilt in the vomer channel.

**Table IV:**
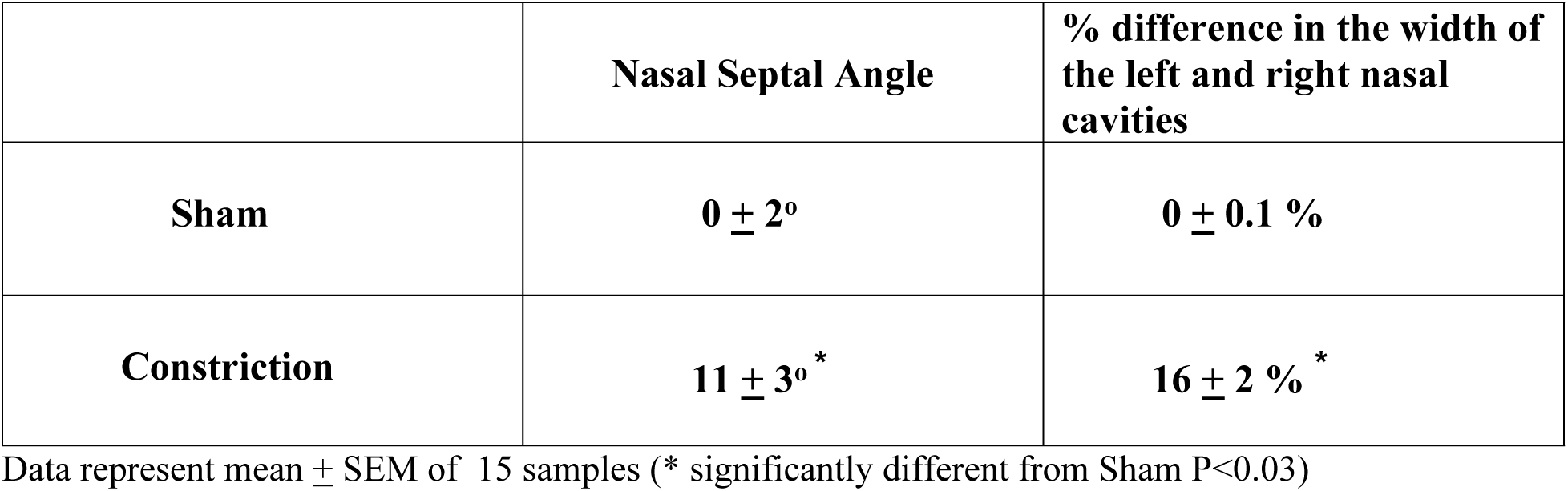
Nasal septal deviation.

|  | Nasal Septal Angle | % difference in the width of the left and right nasal cavities |
| --- | --- | --- |
| Sham | $0 \pm 2^\circ$ | $0 \pm 0.1 \%$ |
| Constriction | $11 \pm 3^\circ$ * | $16 \pm 2 \%$ * |

Data represent mean ± SEM of 15 samples (\* significantly different from Sham $P < 0.03$ )

While in the sham group, the left and right nasal cavities were similar in size, in the experimental group, the size of the nasal cavities was significantly wider on the opposite side of the nasal septal deviation (Table IV) (p<0.03).

The vomer bone, which holds the nasal septum in a channel structure, demonstrated a tilt in the channel. This tilt followed the nasal septum deviation and was more frequent toward the lower part of the nasal septal deviation (Figure 7B).

### Mandibular shift contributed to the nasal floor slanting

To determine whether nasal floor slanting was related to the compression force itself, we reversed the direction of the force and applied tensile force (expansion force). In response to tensile forces, expansion of the dental arches was observed (data not shown). While some animals did not show a shift in the mandible, some shifted to the left, and some shifted to the right, with no significant difference in trend (p>0.05) (Table V). The animals that did not show a mandibular shift did not show slanting of the nasal floor (Figure 8A), while the animals that received tensile forces and developed a mandibular shift developed a mild-to-moderate slanting. The direction of the slant and mandibular shift was the same (Figure 8 B). This observation demonstrated that in addition to constriction of the maxilla, a mandibular shift can also contribute to nasal floor slanting, however, to a lesser degree. Due to mild nasal floor slanting, no septal deviation was observed.

**Figure 8:**
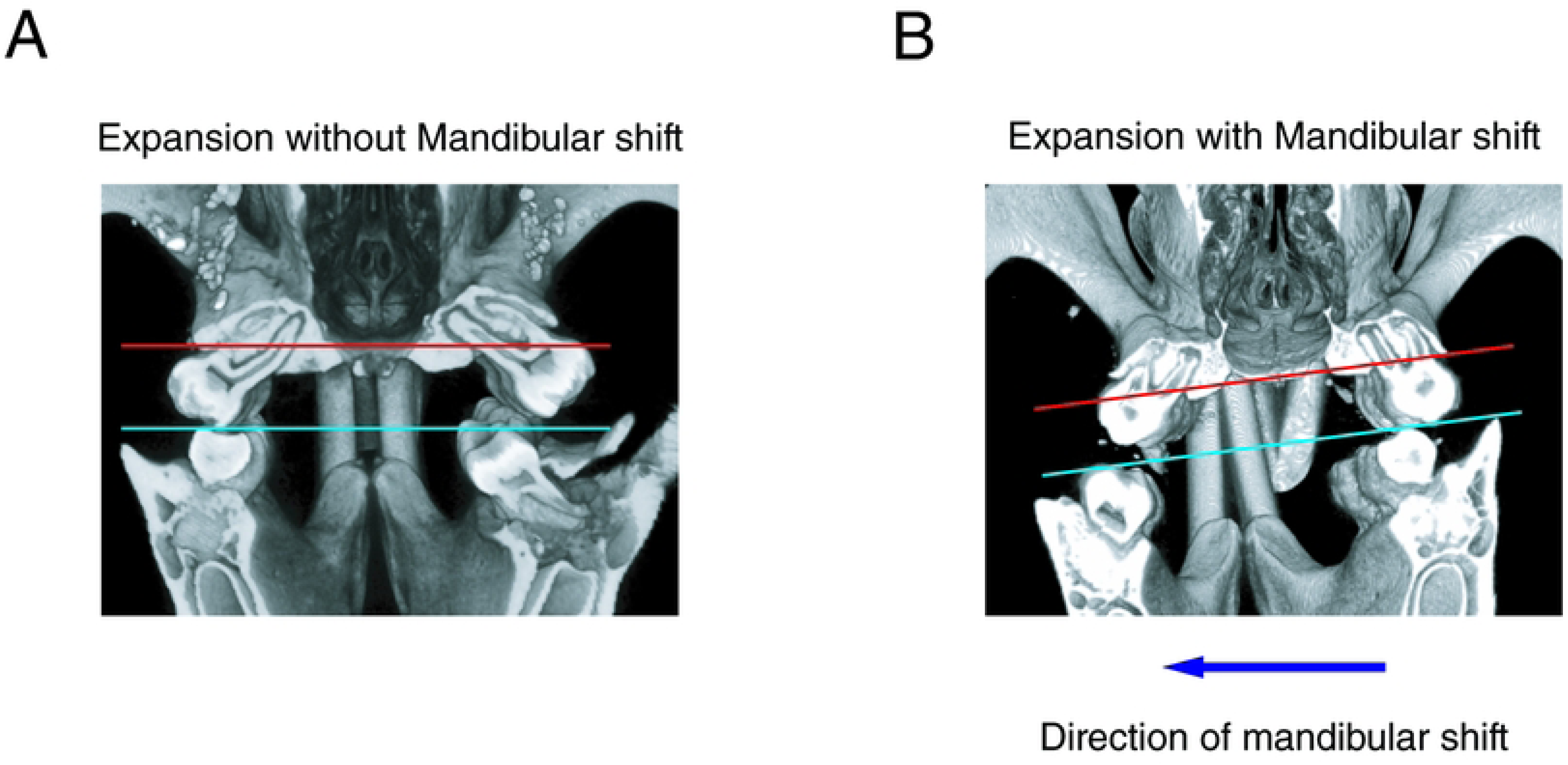
**Tensile forces alone did not increase nasal floor slanting**. **(A)** Animals that did not adopt a mandibular shift did not show any nasal floor slanting. **(B)** Animals that adopted a mandibular shift demonstrated nasal floor slanting, and the direction of slant was similar to the direction of mandibular shift.

**Table V:** Nasal floor slanting in response to expansion.

| Expansion group | Number of animals | Nasal floor slanting |
| --- | --- | --- |
| No mandibular shift | 6 | 0 ± 2 |
| Mandibular shift to the left | 4 | 7 ± 3* |
| Mandibular shift to the right | 5 | 8 ± 3* |

All the nasal floor slants were in the same direction as the direction of mandibular shift. Data represent mean ± SEM of 15 samples (\* significantly different from Sham P<0.05)

## Discussion

Our experiments directly demonstrate that constriction of the maxilla had a significant effect on the nasal septum shape. This agrees with previous findings that found a correlation between maxillary width and nasal septal deviation in humans [42, 43]. This finding has significant clinical importance in light of an increase in the incidence of maxillary constriction in patients prior to orthodontic treatment. The causes of the constriction can be due to a number of environmental factors, including ecological allergens/nasal obstruction, dietary changes, and decreased rates of breastfeeding [28–37]. However, how the maxillary constriction contributes to the development of nasal septum deviation is not clear. Three possibilities exist.

One possible way maxillary constriction may affect the growth of the nasal septum is by creating a growth mismatch between the septum and maxilla. This mismatch may cause the nasal septum (cartilage and bone) to bulge away from the midsagittal plane. This hypothesis aligns with previous studies in humans, which have shown that if normal midline growth of the nasal septum is disrupted, the frequency of septal deviation increases [3, 9, 20, 44, 45].

For this hypothesis to be correct, we need to demonstrate that the nasal septum exhibits robust interstitial growth that persists even in the absence of the space required for growth of the septum due to maxillary constriction. But which part of the nasal septum has the interstitial growth?

The nasal septum is composed of three parts: the cartilaginous septum and the bony septum, which is composed of the perpendicular plate of ethmoid and the vomer. The ossification of the vomer through the intramembranous pathway typically is completed by birth, and the ossification of the perpendicular plate accelerates until the age of ten, then slows down significantly afterward [46, 47]. It has been established that the bony part of the septum does not have interstitial growth and primarily functions as a growth site [48, 49]. On the other hand, since the nasal cartilage is considered the anterior extension of the chondrocranium, it is regarded as a primary cartilage and, therefore, has interstitial growth. It has been suggested that this interstitial growth can produce enough mechanical forces that may help in the maxilla’s displacement and suture activation and, therefore, cause maxillary growth [50, 51]. Thus, the nasal septum has been proposed as the growth center for the maxillary complex [8, 52–59]. While this claim is evident primarily in long-snouted animal models, [60] it is controversial for the human facial skeleton [61]. One group believes that the nasal septum in humans is similar to that of many long-snouted animals and acts as a growth center for the maxilla [6–9, 53, 57, 61], and nasal septal deviation can produce a force that can cause asymmetry in the skull [61–65]. On the other hand, many others could not find the evidence that the mechanical force produced by nasal septum interstitial growth was enough to displace the maxilla. These scientists believe that the soft tissue (periosteal matrix) and oronasal spaces (capsular matrix) are responsible for maxillary growth [48, 66, 67] and nasal septum growth is secondary to midfacial growth. They believe that the nasal septum has more mechanical importance in transferring occlusal forces to the skull than growth importance [61, 68]. The opponents to the role of the nasal septum as a growth center argue that if the mechanical force produced by the nasal septum was large enough, it would prevent the development of nasal septum deviation in the first place by pushing the maxilla downward and forward.

While the interstitial growth of the nasal septum is undeniable, the magnitude of its contribution to nasal septal deviation is questionable. This is because the majority of the nasal septum’s growth decreases significantly by the age of two, while maxillary growth continues by age 14-16 [69]. Based on this observation, nasal septal growth can only partially explain the development of septal deviation.

The second possibility that may explain why maxillary constriction affects the nasal septum deviation is that constriction pushes the palatal shelves upward toward the space occupied by the nasal septum, causing a deviation in the septum. This bulging in response to constriction can be explained mechanically by the appearance of significant moments in the system (Figure 9). We argue that the constriction forces applied far from the center of resistance of each hemimaxilla can produce considerable moments in the system that push the maxilla up in the midline toward the nasal cavity and decrease the space for the septum (Figure 9B). If this hypothesis is correct, one would expect the application of tensile forces to produce opposite moments, and decrease bulging in the middle of the nasal floor, which was observed in our experiment (Figure 10). This analysis is in agreement with previous observations that nasal septal deviation is associated with an increase in palatal arch height [3, 4, 11, 70].

**Figure 9:**
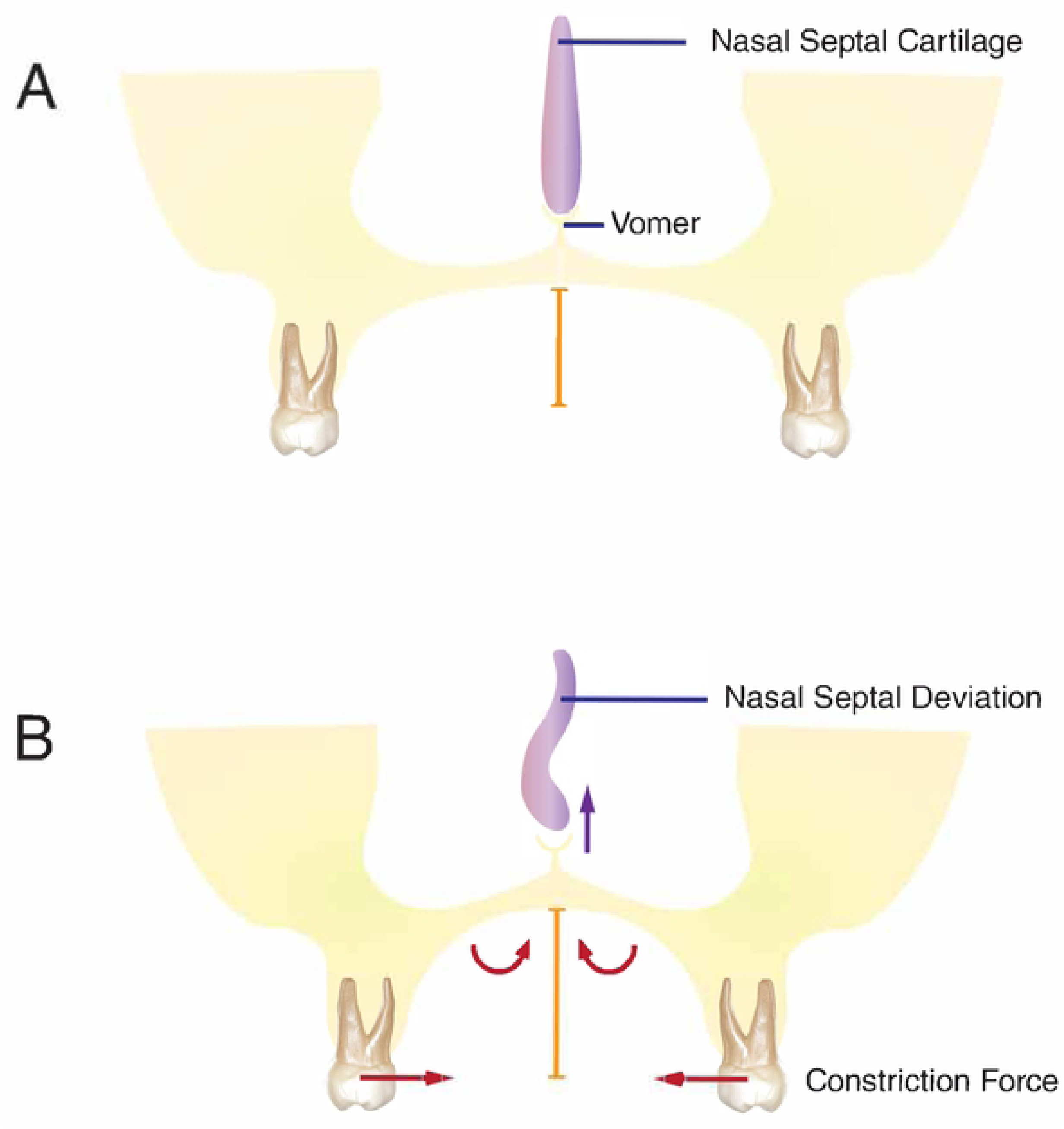
Constriction caused large moments toward the palate. **(A)** In a normal maxilla, the palatal height is not large (orange scale bar), and the palate is wide. No nasal septal deviation is observed in this condition. **(B)** Constricting horizontal forces produced large moments toward the palate (red curved arrows), which pushed both the right and left palatal shelves up and caused bulging in the floor of the nose, increased palatal height (compare orange scale bar with the one in A), and the development of a nasal septal deviation.

**Figure 10:**
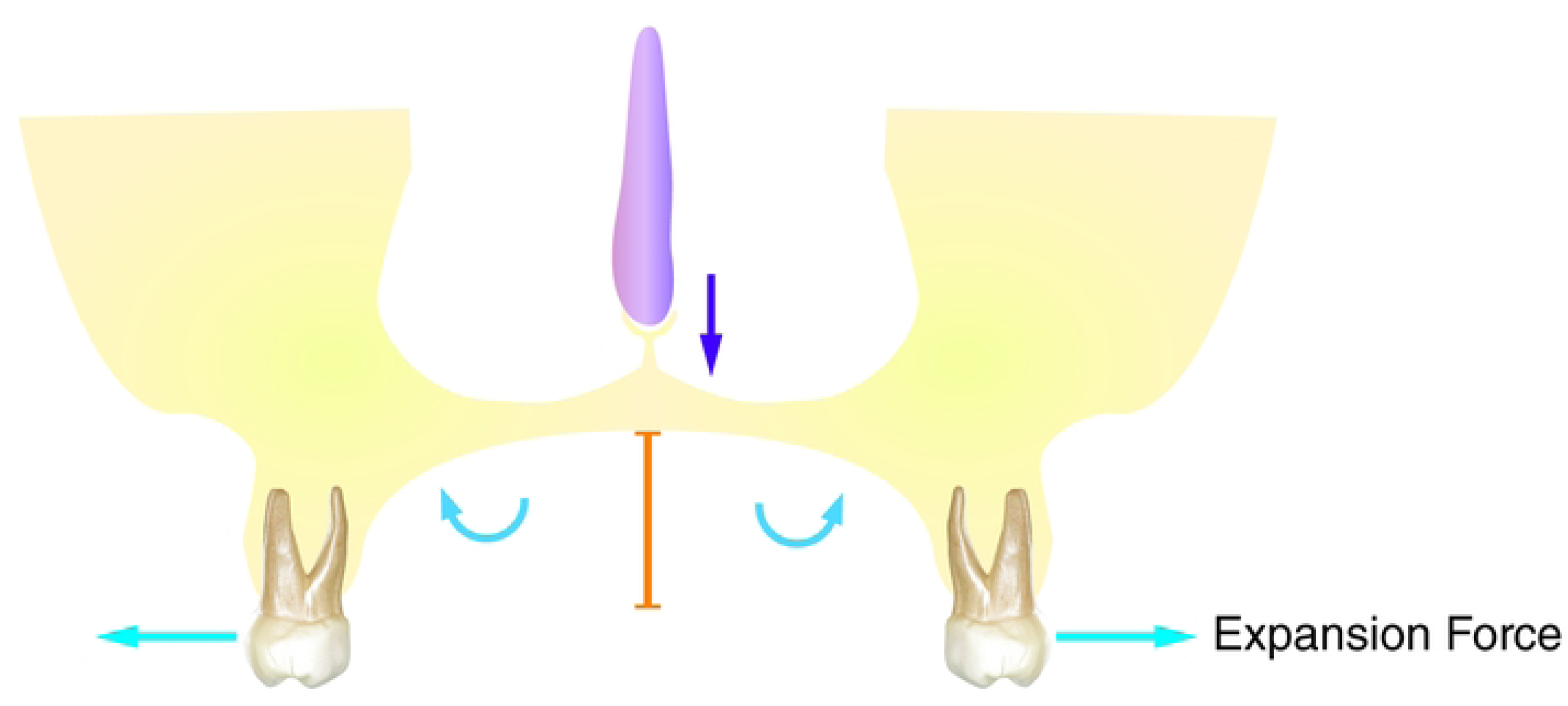
Expansion caused moments toward the buccal plates. In response to expansion forces, horizontal forces appear in the system, which not only expand the maxilla but also produce large moments towards the buccal plates that turn the palatal shelves ventrally. In this condition, no bulging in the floor of the nose and no nasal septal deviation were observed, and the palatal height decreased (vertical orange scale bar).

Symmetrical moments produced by constricting forces should cause bulging in the middle of the nasal floor, but no slanting should occur. However, in our experiment, significant nasal floor slanting was observed, indicating that unequal mechanical factors altered the balance between the right and left sides of the nasal floor. In other words, constricting force alone could not explain the slant in the nasal floor.

Maxillary constriction prevents proper contact between the posterior teeth and forcing the mandible to shift to one side to establish a posterior occlusion, which results in a crossbite, as was observed in our experiments. This produces large horizontal forces toward the medial in the non-crossbite side.

These forces produce an additional moment that can push the palate even further up on that side (Figure 11). On the other hand, on the crossbite side, the horizontal forces remain small and directed laterally, producing a lateral moment that rotates the palate in the opposite direction of the moment created by constricting forces, thereby reducing the magnitude of the rotation of the palatal shelf on that side. This differential moment between the left and right contributes to the slanting of the nasal floor.

**Figure 11:**
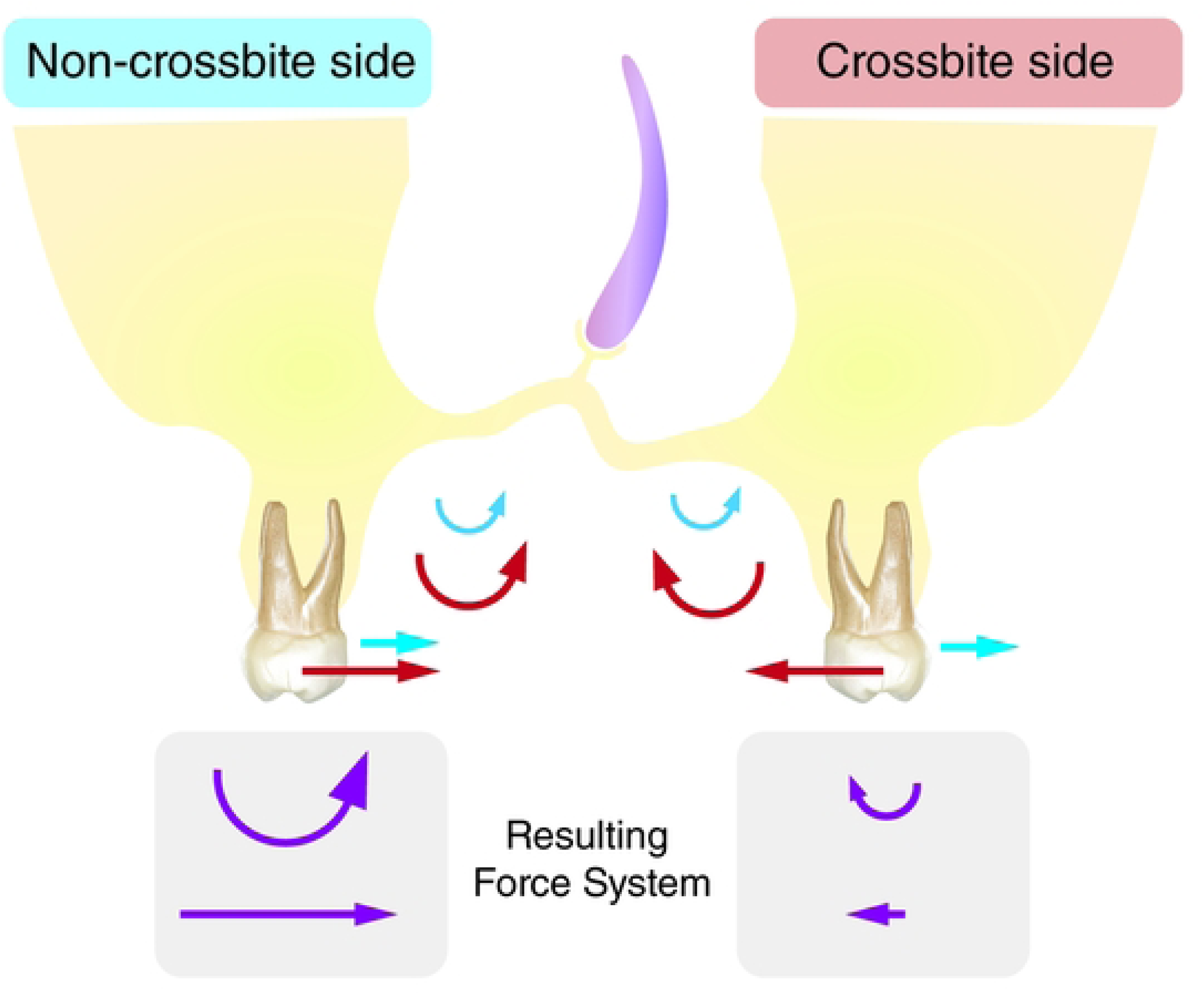
Mandible shift produced additional moments in the system that caused nasal floor slanting. In the presence of constricting forces, the mandible shifted toward one side to produce a more comfortable posterior bite. This resulted in horizontal forces that were medially oriented on the non-crossbite side and laterally oriented on the crossbite side (straight blue arrows). This increased the counterclockwise moment (palatal-oriented moment) in the non-crossbite side and a laterally oriented moment on the crossbite side (curved blue arrows), which can cause significant slanting in the nasal floor. Moments caused by constricting forces are shown as curved red arrows, and resulting moments are shown as curved purple arrows.

In our study, no specific trend was observed in the shift of the mandible toward one side. However, the direction of shift of the mandible and nasal floor cant was always the same. To further test the hypothesis that the shift of the mandible contributes to the slanting of the nasal floor, we studied animals exposed to expansion (tensile forces). Here, we expected no slanting of the nasal floor. However, similar to animals that received the constriction forces, the bite in these animals changed, and the majority of the animals demonstrated a shift in their mandible to develop a normal bite on one side and a scissor bite on the other side (Figure 8). However, some animals were able to continue biting in the center. In none of the animals did the expander cause bulging of the nasal floor as was expected. In the absence of mandibular shift, no slanting of the nasal floor was observed. But in the presence of mandibular shift, mild to moderate nasal floor slanting was observed. In this scenario, the direction of nasal floor slanting and mandibular shift was the same. These experiments together demonstrate that the slanting was caused by mandibular shift and could be exaggerated in the presence of constricting forces. It should be emphasized that the shift itself did not cause the large horizontal forces and moments. The shift is associated with a change in biomechanics of occlusion, such as a change in the inclination of the lower molars, that will continue even when the shift of the mandible gradually changes to adaptation of form and development of permanent asymmetry. It should also be noted that since the changes in nasal septal deviation occur all throughout life [9, 46, 71, 72], one can expect the dynamic changes in occlusion that can occur throughout life to contribute to changes in nasal deviation.

We know that the nasal floor is part of the palatal process of the maxilla, and its growth is a combination of bony resorption on the nasal side, along with bony apposition over the oral surface, which is considered part of natural cortical drift [49]. The change in loading of the maxilla and the appearance of moments and horizontal forces due to constriction forces and mandibular shift produce cortical drifting in the opposite direction, which causes changes in the floor of the nose and the height of the palate [73, 74]. We believe that this change in occlusal forces and the appearance of new moments that caused nasal floor slanting can be the third source for the development of nasal deviation. Nasal floor slanting is accompanied by a change in the direction of the vomer, as was observed in our experiment here. This is important, since the vomer holds the base of the nasal septal cartilage (Figure 12). Nasal septal cartilage lies in the vomerine groove, a shallow channel at the superior border of the vomer bone. Therefore, a change in the direction of the vomer could easily cause nasal septal deviation. Our observation that nasal floor slanting and vomer inclination could be considered as a third factor in the development of nasal septal deviation is in agreement with previous studies, which show a high incidence of nasal septal deviation associated with nasal floor slanting [65, 69, 75].

**Figure 12.**
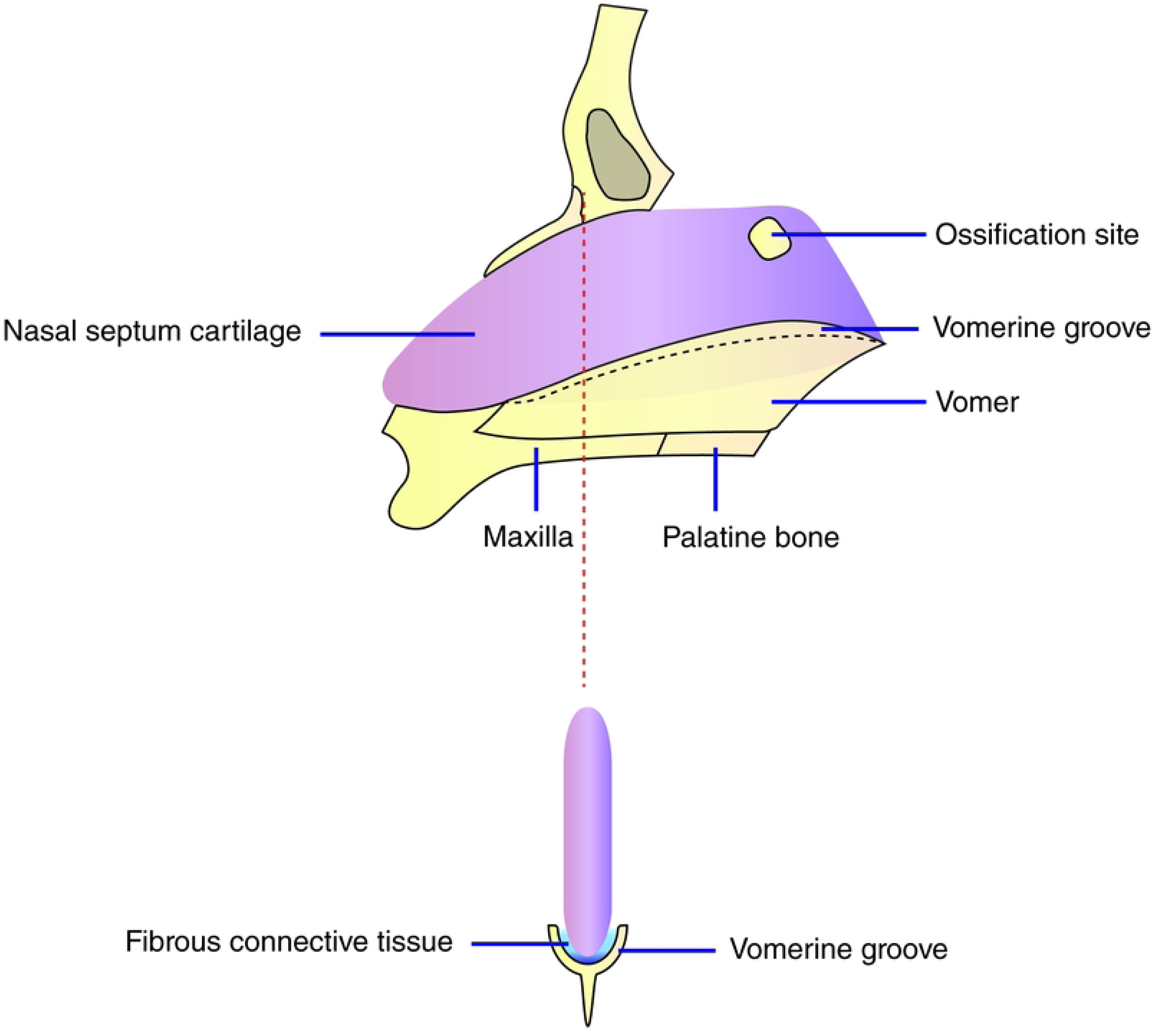
Schematic view of the vomerine groove. (**A**) Sagittal view of cartilage at the embryonic stage demonstrates that cartilage at the inferior border is located in the channel at the superior border of the vomer bone (vomerine groove). The black dashed line reflects the extension of cartilage between the perpendicular plate and vomer bone (vomerine process). (**B**) A cross-section of the nasal septum (frontal view) along the red dashed line from panel A demonstrates how the inferior border of cartilage is embraced by the vomer bone. Fibrous tissue connects these two structures.

In our experiments, we did not find a significant difference in the direction of the vomer nor nasal septal deviation toward one side, which suggests that there is a complex mechanical environment in the oral and nasal cavities. This differs from previous studies, which noted that nasal septum deviation is more pronounced on the lower side of the nasal floor [3, 43, 75, 76]. This difference may be due to the limited sample size in our experiment.

In our experiments, constriction of the maxilla was accompanied by C-shape deformation of the nasal septum, in accordance with the classification system proposed by Guyuron [77]. However, we did not observe any other type of deformation. This can be due to a short duration of study. Our observation is in agreement with previous reports that have shown the most frequent nasal septal deviation associated with nasal floor slanting is the C-shape deformity [43]. Perhaps that can explain why the application of expansion forces can be more successful in the treatment of C-shape deviation, as has been reported before [78].

This study shed light on nasal floor slanting as the third source of nasal septal deviation. However, in addition to nasal septal deviation, nasal slanting has other side effects. Studies demonstrated that severe slanting of the nasal floor may also contribute to nasal obstruction [79], which emphasizes the importance of clinical diagnosis of constricted maxilla that can cause nasal floor slanting.

Rats in this experiment demonstrated clockwise rotation of the mandible, increased lower facial height, and decreased posterior facial height. These changes were not due to a change in the animal’s breathing pattern and were induced mostly by constriction of the upper arch and uprighting of the upper posterior teeth. However, they produced similar skeletal effects to those observed in mouth-breathing patients [80, 81]. This similarity can be due to the effect mouth breathing has on the constriction of the upper arch [70]. Chronic mouth breathing can increase the constricting forces on the maxilla, which will affect transverse and vertical growth of the maxilla [11–13].

It should be emphasized that our study did not test nor deny the possible effect of septal growth on the maxilla [51, 54–56]. However, we demonstrated that even in the presence of the mechanical forces from the septum, maxillary growth can easily be derailed by many other factors. In fact, the maxilla itself can change the direction of septal growth, which can cause chronic blockage of the nasal cavity and mouth breathing [36, 82–84]. Based on this reciprocal theory, deformity of craniofacial structures worsens nasal obstruction and chronic mouth breathing, which, in turn, leads to craniofacial maldevelopment. This negative feedback loop emphasizes the need to interrupt the cycle as early as possible through proper diagnosis and early interceptive treatment.

## Conclusion

This article, from many aspects, is original and innovative. First, it sheds light on maxillary constriction as one of the leading developmental factors in nasal septal deviation. Second, this study for the first time emphasizes that the nasal cavity and oral cavity as two systems that have significant reciprocal effect on each other’s form. Third, it identified two possible pathways by which maxillary constriction can affect nasal septal deviation: on one pathway, maxillary constriction creates mechanical forces and moments that cause bulging of nasal floor, which invades the space for the septum; and on the other pathway maxillary constriction can change occlusal forces, which may lead to nasal floor slanting, vomer deviation, and, therefore, nasal septum deviation. Finally, this article demonstrates how maxillary constriction automatically changes the skeletal form and produces malformities. These findings underscore the importance of recognizing maxillary constriction as a public health issue that can significantly compromise the health of children and adults, necessitating early diagnosis and treatment. Children and pre-pubertal individuals show greater adaptability of the nasal septum following orthopedic interventions, such as maxillary expansion [85]. This is in addition to the other beneficial effects of palatal expansion on increasing the nasal cavity and decreasing inspiratory nasal resistance [86, 87]. Based on this study, there is a significant need for additional research in this field and closer collaboration among different disciplines to address what is becoming a large public health issue.

## Funding Source

This research did not receive any grant from funding agencies in the public, commercial, or not-for-profit sectors. This project was self-funded by CTOR (Hoboken, NJ) .

## Acknowledgments

We acknowledge the Unidad de Modelos Biológicos at the Instituto de Investigaciones Biomédicas, Universidad Nacional Autónoma de México (UNAM), where the animal experiments were performed. Special thanks to DVM. Rubi E. Zavala Gaytán and DVM. Filipo C. Paczka García for their technical support and animal care at the Unidad de Modelos Biológicos

## Notes

### Competing Interest Statement

The authors have declared no competing interest.

